# Declining muscle hyperplasia in juvenile trout is driven by rapid limitation of muscle stem cell capacity and niche functionality

**DOI:** 10.64898/2026.01.28.702281

**Authors:** Sabrina Jagot, Nathalie Sabin, Cécile Rallière, Adèle Branthonne, Morgane Chesnais, Cécile Duret, Jérôme Bugeon, Pierre-Yves Rescan, Karl Rouger, Jean-Charles Gabillard

**Affiliations:** INRAE, LPGP, 35000, Rennes, France; Oniris, INRAE, PAnTher, 44307, Nantes, France

## Abstract

Unlike mammals and birds, where new muscle fiber formation (hyperplasia) ceases around birth, large and fast-growing fish such as rainbow trout undergo a spectacular post-hatching surge of hyperplasia, followed by a considerably delayed hyperplasia decline. This study investigated the roles of the satellite cells (SCs) and their niche in this decline by determining the number and the myogenic capacity of the muscle progenitors as well as the functionality of their direct tissue environment. Histological analysis revealed a significant decrease in hyperplasia (fibers <25 µm) and SC numbers (Pax7^+^) between 10 g and 500 g trout. Transplantation experiments using muscle-derived cells (MDCs) from *mlc2*-GFP transgenic trout (10 g to 2 kg donors into 10 g to 2 kg recipients) demonstrated a marked decline in both intrinsic myogenic capacity and niche functionality as trout grow from 10 g to 500 g. Detailed analyses of GFP^+^ fibers produced after transplantation showed an enrichment of small-diameter GFP^+^ fibers in 10 g but not 100 g trout recipient muscles, showing a rapid impairment in niche ability to support hyperplasia. In addition, transplantation of MDCs from trout of different ages but the same weight, showed that increasing trout weight, but not aging, is associated with an impairment of the myogenic capacity of progenitors and their niche. Overall, these findings show that the muscle hyperplasia decline in trout is primarily driven by early impairment of the SC niche, followed by a reduction in their myogenic capacity and number, with weight gain playing a more critical role than aging.

## Introduction

In vertebrates, skeletal muscle is primarily made up of multinucleated muscle fibers, formed mainly during embryonic and fetal development (1). After birth, the formation of new fibers (hyperplasia) no longer exists in mouse (2), pig (3), chickens (4), or persists for only a few weeks in rats (5). In stark contrast, in large and fast-growing fish such as salmonids, a spectacular surge of hyperplasia occurs during juvenile muscle growth, as illustrated by the 200-fold increase in fiber number between hatching and a 4 kg salmon (6). The hyperplasia decline is also present in these fish but considerably delayed, as the number of muscle fibers is considered to cease when the body length reaches about half its maximum (7).

Post-natal muscle hyperplasia requires an active contribution of muscle stem cells also called satellite cells (SCs), which are located between the basal and plasma membranes, defining their anatomical niche (8, 9). SCs are commonly identified using the well-known Pax7 marker, essential for cell survival, cell fate and self-renewal (10). Upon activation in the context of growth or injury, these cells generate myogenic precursors that either fuse to existing muscle fibers to increase their size (hypertrophy) or fuse together to form new muscle fibers (hyperplasia) (11). With age, exhaustion of SCs strongly compromise the muscle regenerative capacity (12–14). Similar results have been observed in the context of diseases such as muscular dystrophies (12). Experimentally, transplantation of SCs from young mice produces better muscle regeneration than that with adult SCs, reinforcing the hypothesis of a reduction in the intrinsic myogenic capacity of SCs with aging (15). Concomitantly, the age of the recipient has been shown to be critical for efficient muscle regeneration, highlighting that the tissue context, *i.e*., muscle niche, is also essential to promote new fiber formation (16).

Whereas these hallmarks of reduced hyperplasia capacity are evidenced in aged muscle of mammals, the hyperplasia decline in trout is considered to occur during the juvenile stage (17, 18). The juvenile period is characterized by an early juvenile phase from 5 g to around 500 g (∼12 month) followed by a late juvenile phase until sexual maturation (1.5 - 2 kg; 2-3 years). We have previously observed a lower proportion of proliferative SCs in 500 g than in 2 g trout and a quiescent state of muscle-derived cells (MDCs) in 2 kg trout. Moreover, only a few myofibers were produced after muscle injury in 1.5 kg trout, and the ability of the muscle to regenerate appeared to be impaired, as suggested by the strong development of connective tissue at the injury site (19). Thus, the myogenic capacity of the trout muscle seems to switch from high to low levels during the juvenile period.

In the present study, we followed the evolution of hyperplasia during juvenile period of rainbow trout to precisely define its end. The aim of our work was to determine the myogenic capacity of SCs and the functionality of their niche in order to define their respective roles in the decline of muscle hyperplasia in trout.

## Results

### Muscle hyperplasia decreases sharply in early juvenile trout stage

To accurately determine the kinetics of muscle hyperplasia, the diameter of fibers was measured in the white muscle of juvenile trout from early (10 g) to late (2 kg) juvenile stages. Cross-sectional images of WGA-labeled muscle continuously showed the simultaneous presence of small (<25 µm) and large (>100 µm) fibers, giving the trout muscle a mosaic appearance (Fig. 1*A*). Fiber distribution density showed a clearly difference in muscle fiber content as a function of fish weight, with maximum diameters of 100 µm for the 10 g trout and 220 µm for the 2 kg immature trout (Fig. 1*B*). Furthermore, the highest densities were found for 25-30 µm, 40-50 µm and 60-70 µm fibers for 10 g, 100 g and 500 g trout, respectively. In addition, a low proportion of small fibers was noted in trout weighing over 500 g. To further characterize the kinetics of new fiber formation, we focused on fibers smaller than 25 µm in diameter as an indicator of muscle hyperplasia activity. Consistent with fiber distribution density, 34±4% and 11±8% of <25 µm fibers were presents in 10 g and 500 g trout muscle respectively, attesting to the sharp decrease (p<0.01; Fig. 1*C*) of muscle hyperplasia. Thereafter, the proportion of <25 µm fibers, corresponding to less than 6%, remained within the same range between 1 kg, 1.5 kg and 2 kg trout. To take better account of the huge difference in fiber diameter, the proportion of surface area occupied by the <25 µm fiber over the entire area was determined. It represented 4.7 %, 0.7 % and less than 0.3 % in 10 g, 500 g (p<0.001) and larger trout, respectively (Fig. 1*D*). Taken together, these results showed that muscle hyperplasia in the white muscle decreases sharply and rapidly in early juvenile trout stage between 10 g and 500 g and reaches a minimum value from 1 kg.

**Figure 1:**
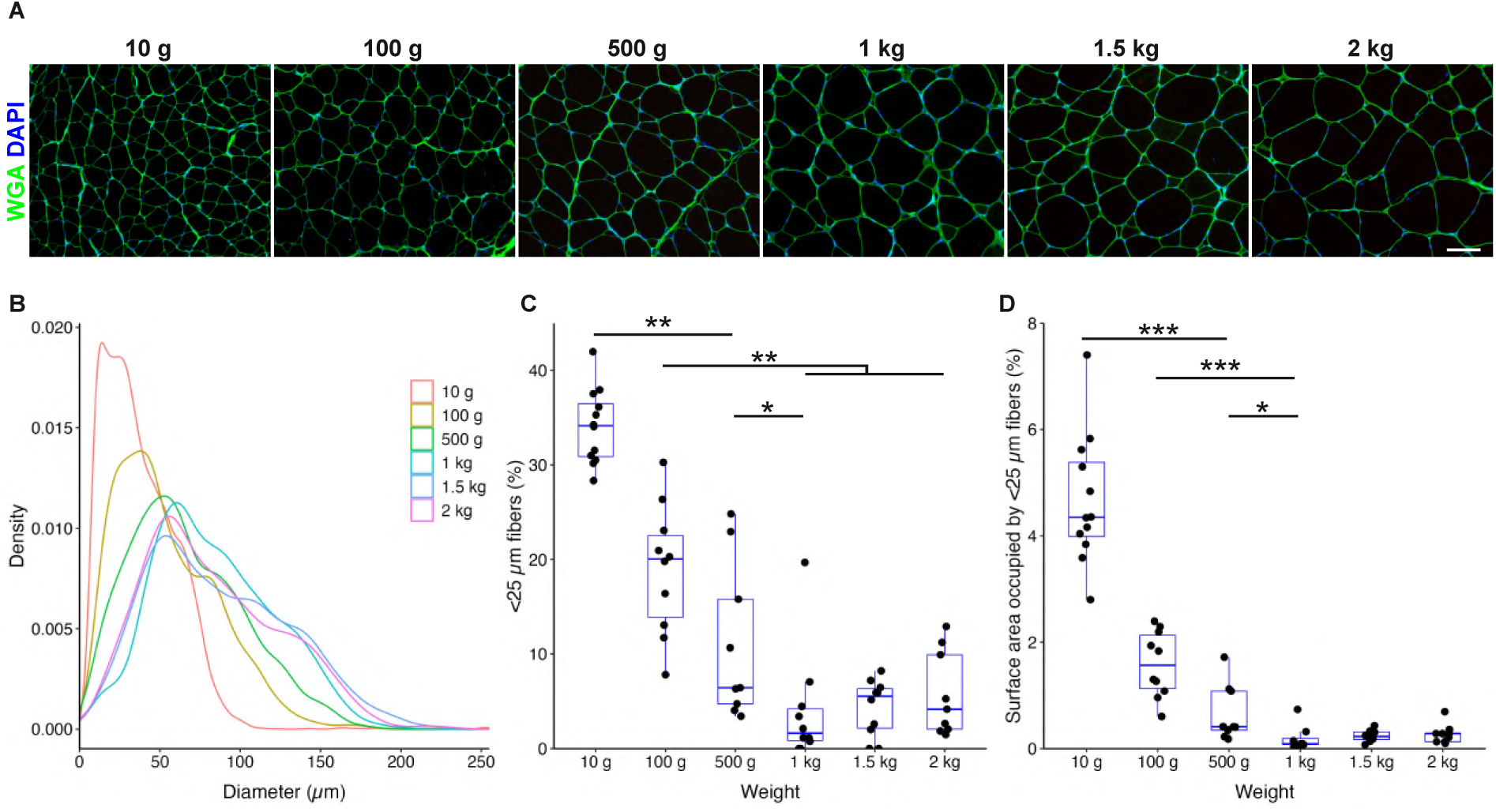
Muscle hyperplasia in trout decreases sharply in juvenile trout. (A) Cross-sections of white muscle from 10 g to 2 kg trout were stained with Alexa 488-conjugated wheat germ agglutinin (WGA; green) to visualize the extracellular matrix. The nuclei are counterstained with DAPI and the scale bar corresponds to 100 µm. (B) Density function of fiber diameter (µm) in white muscle from 10 g to 2 kg trout. (C) Percentage of myofibers with a diameter of less than 25 µm. (D) Percentage of the surface occupied by the fibers of less than 25 µm in diameter. The data in (C) and (D) are presented as a boxplot (N = 8-10). All statistical significances were calculated by the non-parametric Kruskal–Wallis rank test followed by the post-hoc Dunn test. Significance levels: *p<0.05, **p<0.01, ***p<0.001.

### Number of pax7^+^ cells decreases sharply in muscle at early juvenile trout stage

We recently used a highly sensitive *in situ* hybridization method to accurately detect the expression of *pax7,* the canonical SC marker (20). Here, this method enabled us to visualize the SCs in their typical position under the basal membrane (Fig. 2*A*) and to count 23.8±3.0 and 2.3±1.5 SCs /mm^2^ in muscle section of 10 g and 500 g trout, respectively (Fig. 2*B*). The density of SCs is 10 times higher in the muscle of 10 g trout compared with that of 500 g trout (p<0.001). Thereafter, SC density reached a minimum plateau and remained comparable up to 2 kg trout. As increasing fiber size with fish weight could induce a decrease in SC density, the number of *pax7*^+^ cells /fiber cross-section was then determined (Fig. 2*C*). The evolution of the number of *pax7*^+^ cells was the same as described above. Finally, the number of *pax7*^+^ cells was related to the total number of nuclei counted in a cross-section image of white muscle. Again, the results clearly showed a 10-fold reduction (3.0±0.4 *versus* 0.4±0.3; p<0.001) in the proportion of *pax7*^+^ cells between 10 g and 500 g juvenile trout (Fig. 2*D*). Collectively, these data show a dynamic evolution in the number of SCs, which falls during the early juvenile stage of trout, leading to a low value in trout over 500 g.

**Figure 2:**
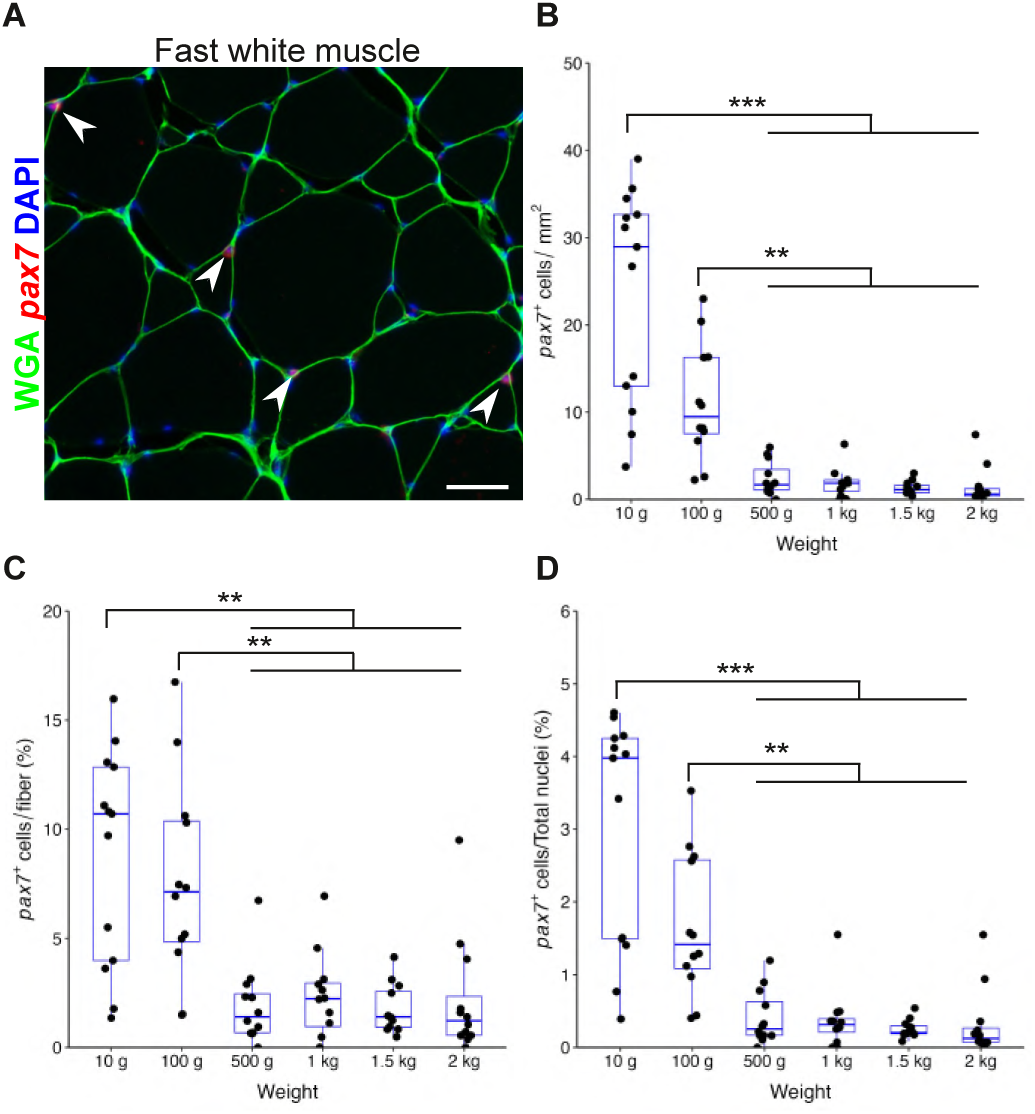
The density of *pax7^+^* cells in white muscle decreases sharply in early juvenile trout. (A) Cross-sections of trout white muscle were analyzed by *in situ* hybridization for *pax7* (red) and the extracellular matrix was stained with Alexa 488-conjugated wheat germ agglutinin (WGA; green). Arrowheads indicate *pax7^+^* cells in the white muscle. The nuclei are counterstained with DAPI and the scale bar corresponds to 100 µm. (B) Quantification of the *pax7^+^* cells /mm^2^ in white muscle from 10 g to 2 kg trout. (C) Quantification of the percentage of *pax7^+^* cells /myofiber in white muscle from 10 g to 2 kg trout. (D) Quantification of the percentage of *pax7^+^* cells /total nuclei in white muscle from 10 g to 2 kg trout. The data in (B), (C) and (D) are presented as a boxplot (N = 8-10). All statistical significances were calculated by the non-parametric Kruskal–Wallis rank test followed by the post-hoc Dunn test. Significance levels: *p<0.05, **p<0.01, ***p<0.001.

### Increasing donor and recipient weight impairs the ability of myogenic progenitors to differentiate

Given the highly evolutionary nature of the SC number and the shift from hyperplastic to hypertrophic growth during the juvenile period, we wanted to further explore the impact of trout weight on both the myogenic progenitor capacity and the muscle niche functionality. To that end, transplantation assays were performed by using MDCs extracted from *mlc2*-GFP transgenic trout line as donor, which allows us to specifically detect differentiated myogenic progenitors thanks to exclusive GFP expression (21). This was supported by preliminary work in wild-type (WT) recipient showing that the GFP signal increased up to 29 days post-transplantation and remained stable at least up to 104 days (Supplemental Figure S1). Also, very few *CD8*^+^ T-cells were identified at the transplantation site in 10 g and 1 kg trout, evoking an absence of a rejection mechanism (Supplemental Figure S2). First, to assess the niche ability to support new fiber formation, *mlc2*-GFP MDCs of donor 10 g trout were transplanted into early to late juvenile recipient trout (10 g to 2 kg). After 21 days, a strong GFP signal throughout the transplantation area was observed in muscle of 10 g trout (Fig. 3*A, B*). A clear GFP signal was also observed into 100 g trout muscle but over a smaller surface. Subsequently, very weak GFP signal was detected in the white muscle of 500 g to 2 kg trout. Muscles from 10 g and 100 g trout were characterized by a significant signal (GFP/Red surface ratio) compared to heavier trout muscles (p<0.001). These data showed that the tissue environment is less and less supportive of the SC action with weight gain, and that very quickly, *i.e*. from 500 g, it is almost no longer functional.

**Figure 3:**
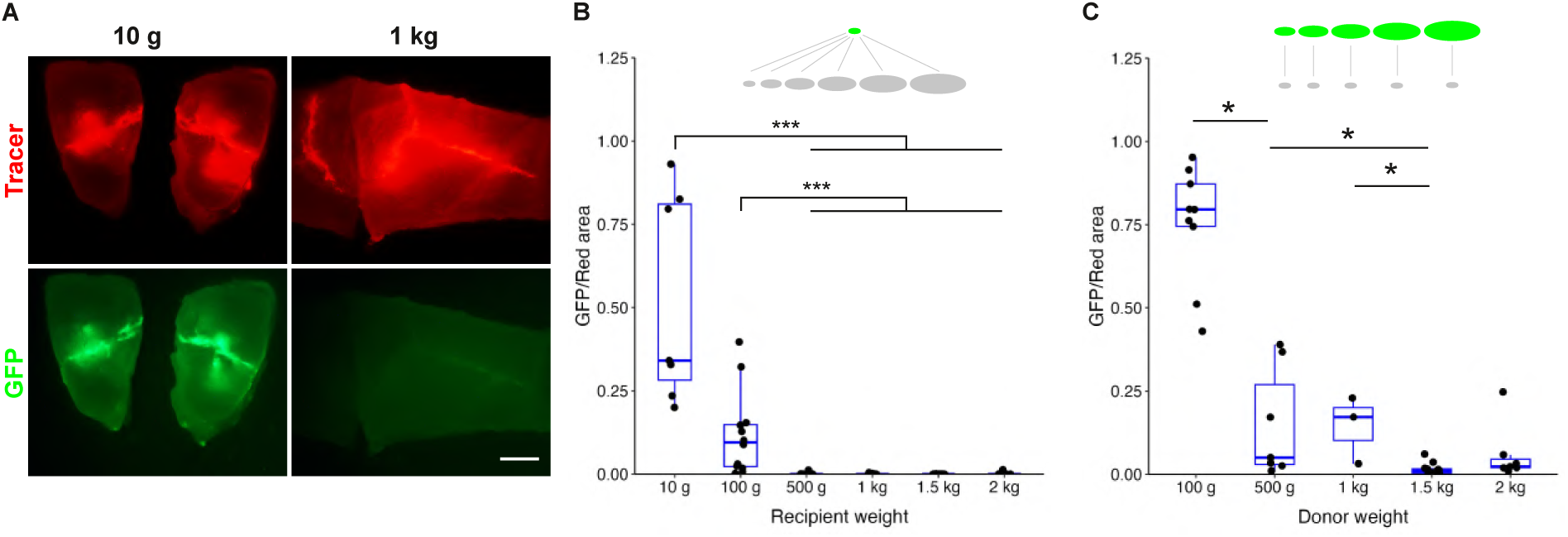
Increasing donor and recipient weight impairs the ability of myogenic progenitors to differentiate. (A) Pictures of muscle samples of 10 g and 1 kg trout, 21 days after transplantation of 1×10^5^ *mlc2*-GFP muscle-derived cells (MDCs) that show the injection site (red tracer) and the endogenous GFP signal (green) resulting from the differentiation of transplanted myogenic progenitors. The scale bar corresponds to 2 mm. (B) Calculation of the ratio between the GFP signal surface and the red surface obtained after transplantation of *mlc2*-GFP MDCs from 10 g trout into muscle of 10 g to 2 kg trout (N = 7-12). (C) Calculation of the ratio between the GFP signal surface and the red surface obtained after transplantation of *mlc2*-GFP MDCs from 10 g to 2 kg trout into muscle of 10 g trout (N = 7-9, except for 1 kg where only 3 fish could be analyzed). The data in (B) and (C) are presented as a boxplot. All statistical significances were calculated by the non-parametric Kruskal–Wallis rank test followed by the post-hoc Dunn test. Significance levels: *p<0.05, ***p<0.001.

Next, to determine whether the intrinsic capacity of SCs changes in relation to fish weight, mlc2-GFP MDCs from early to late juvenile donor trout (100 g to 2 kg) were transplanted into 10 g WT recipient trout. Twenty-one days later, transplanted MDCs from 100 g donor trout gave a strong GFP signal (Fig. 3C). In comparison, the GFP signal was lower with 500 g donor trout (p<0.05) and almost no longer detectable from 1.5 kg showing that the intrinsic capacity of myogenic progenitors is also strongly linked to weight gain, with a marked reduction rapidly observed. Taken together, these results show that the hyperplasic capacity of muscle is particularly strong in early juvenile trout stage, but also rapidly limited by the intrinsic capacities of myogenic precursors and the muscle niche, which both are evolving unfavorably. In line with these observations, muscle regeneration capacity assessed by the number of small fibers formed after mechanical injury, was determined as lower in 1 kg trout than in 10 g trout (P<0.001; Supplemental Figure S3).

### The decline in niche functionality impairs muscle hyperplasia capacity before the intrinsic capacity of myogenic precursors is reduced

Knowing that transplanted SCs generate myogenic progenitors capable of differentiation into recipient muscle after 21 days, we sought to determine in what manner they could participate to hyperplasia or hypertrophy. For that purpose, we determined histologically the size distribution of the GFP^+^ fibers newly formed after 21 days. Muscle cross-sections of 10 g recipient transplanted with mlc2-GFP MDCs extracted from 10 g and 100 g trout were defined by the presence of numerous GFP^+^ fibers (Fig. 4*A*, *B*). An enrichment of GFP^+^ fibers in 10-30 μm diameter range was observed, consistent with the global distribution of the endogenous fiber diameters (Fig. 4*D*, *E*). In return, when MDCs from 10 g trout were transplanted in the muscle of 100 g trout, only a few GFP^+^ fibers were detected (Fig. 4*C*) with a broad size distribution (Fig. 4*F*). These data indicate that the capacities of myogenic progenitors for hyperplasia are present at the first stages of growth and that the muscle niche acts negatively very early on the expression of these intrinsic capacities in favor of an orientation towards a hypertrophic modality.

**Figure 4:**
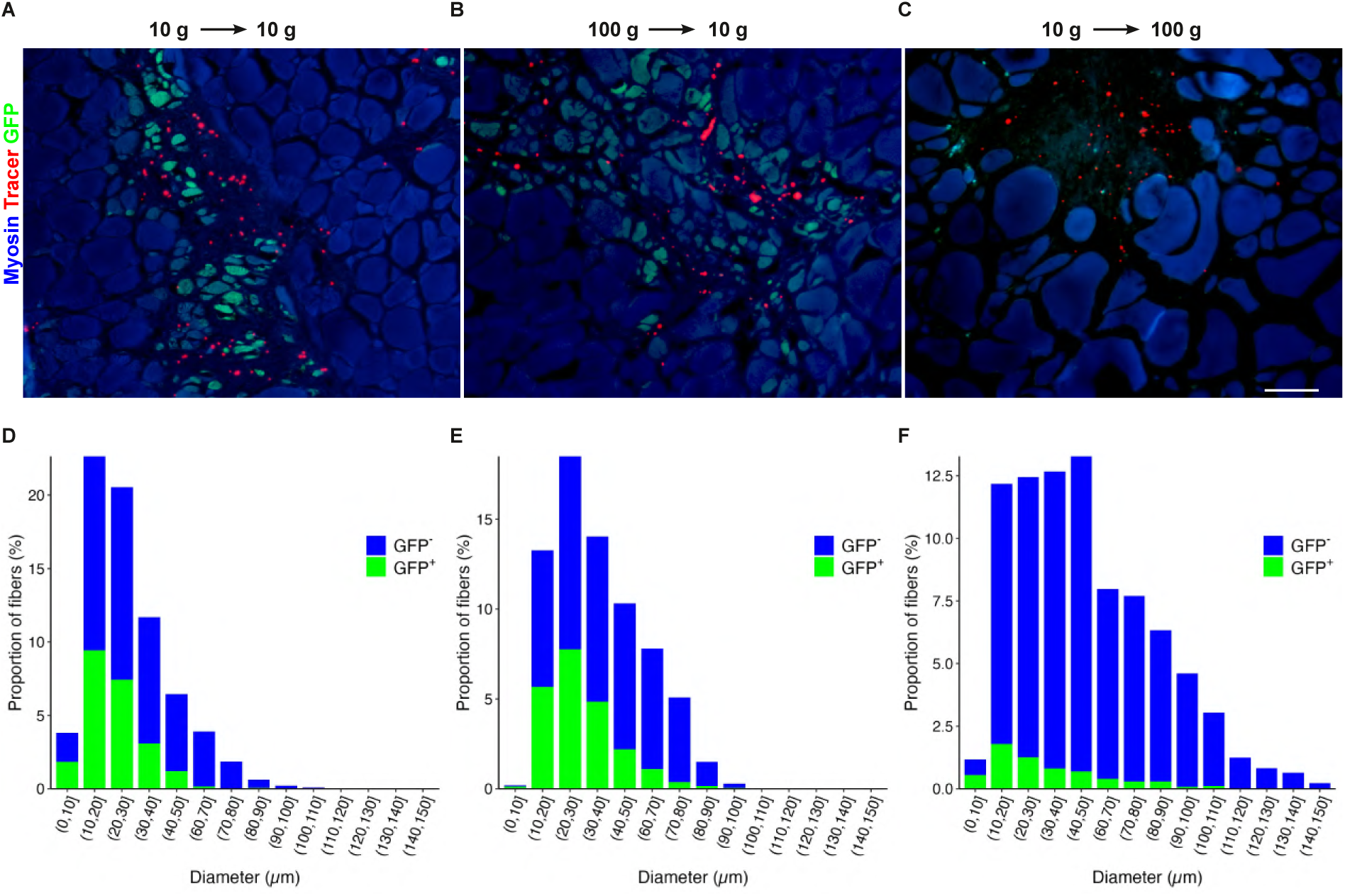
Early decline of muscle hyperplasia capacity in trout. Pictures of muscle cross-sections showing immunolabeling for myosin (blue) endogenous GFP (green) and tracer (red) 21 days after transplantation (A, B, C). The scale bar corresponds to 100 µm. Panels A and D correspond to transplantation of 100 g *mlc2*-GFP muscle-derived cells (MDCs) into muscle of 10 g trout. Panels B and E correspond to transplantation of 10 g *mlc2*-GFP MDCs into muscle of 10 g trout. Panels C and F correspond to transplantation of 10 g *mlc2*-GFP MDCs into muscle of 100 g trout. Panels D, E and F correspond to GFP^+^ and GFP^−^ fiber size distribution.

### Capacity of myogenic progenitors and functionality of their niche decrease with trout weight but not with aging

Considering that trout show exponential growth associated with hyperplastic activity only in the early juvenile stage, we wondered whether increasing weight instead of age might be responsible for the decline in myogenic capacity. To test this hypothesis, we took advantage of the extraordinary plasticity of the trout continuous growth to obtain 20 g and 1 kg fish at the same age (12 months [mo]) by limiting growth through dietary restriction (Fig. 5*A*). We then applied transplantation assay to assess the myogenic capacity of these fish. Transplantation of MDCs extracted from 20 g/6 mo trout into 20 g/6 mo and 20 g/12 mo trout resulted in comparable GFP/Red area ratio of 0.23±0.12 and 0.15±0.14, respectively (Fig. 5*B*). This indicates that the age of the recipient had no major impact on the intrinsic capacity of transplanted progenitors. On the other hand, the GFP/Red ratio observed in 1 kg/12 mo trout receiving MDCs from 20 g/6 mo donors was very limited (0.007±0.014), illustrating a significantly lower myogenic capacity and pointing to a major effect of the weight status of the recipient on the SCs behavior (p<0.01). GFP signals of 0.21±0.24 and 0.007±0.010 were calculated in 20 g/6 mo trout after transplantation of 20 g/12 mo and 1 kg/12 mo trout SCs, respectively (Fig. 5*B*). These values show a very reduced SC myogenic capacity of large donors, indicating a negative effect of donor weight gain on their SC myogenic capacity (p<0.01). Finally, transplantation of MDCs from 20 g/6 mo and 20 g/12 mo trout gave equivalent ratios of 0.23±0.12 and 0.21±0.24 respectively in recipients of 20 g/6 mo, showing no effect of the age of the recipient (Fig. 5*B*). Taken together, these data demonstrate that the myogenic behavior of SCs is conditioned by the weight of both donor and recipient and not by their age, in other words by both the residual intrinsic capacities of the SCs and the signals they receive from the muscle niche.

**Figure 5:**
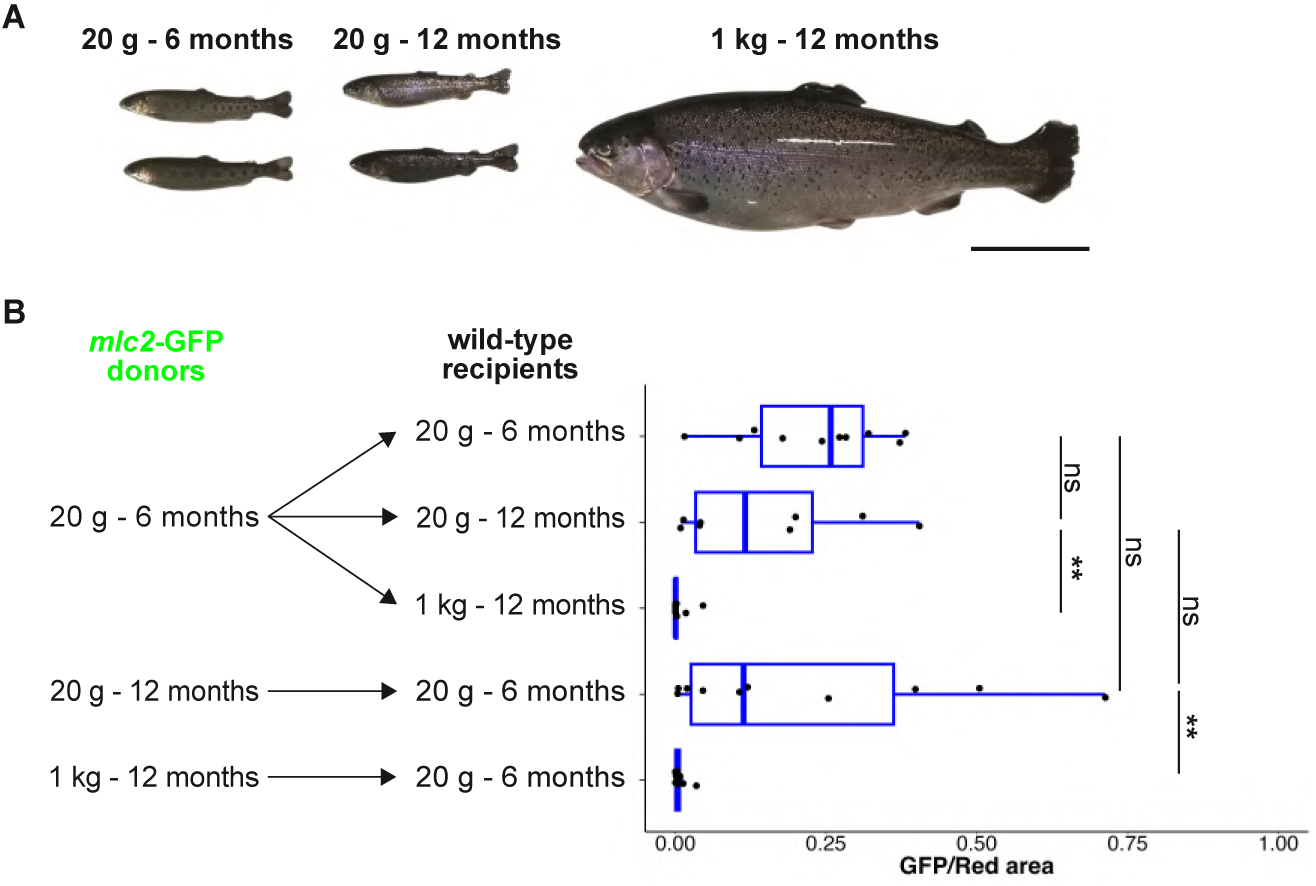
The decline in myogenic capacity of trout muscles is not due to aging but to an increase in weight. (A) Picture of trout for the different combinations of weight/age. The scale bar corresponds to 10 cm. (B) Experimental design of transplantation experiments with different combinations of weight/age trout and calculation of the ratio between the GFP signal surface and the red surface obtained after transplantation of *mlc2*-GFP MDCs into muscle of wild-type trout (N = 8-10). All statistical significances were calculated by the non-parametric Kruskal–Wallis rank test followed by the post-hoc Dunn test. Significance levels: **p<0.01.

## Discussion

The understanding of the muscle hyperplasia process is crucial for deciphering the precise ways in which muscle tissue is formed and to identify the key regulators of SCs during myogenesis and muscle repair. These elements are essential with a view to modulating tissue growth and proposing optimized solutions in the context of cell therapies. The availability of a natural biological model such as trout, which exhibits a spectacular post-hatch surge of muscle hyperplasia, offers a unique study context in vertebrates. Here, we sought to define the quantitative and qualitative characteristics of SCs during juvenile growth in terms of myogenicity, but also to look at the role played by the muscle niche on their potential. The searched aim was to shed light on the reasons for the decline in muscle hyperplasia.

In 1988, a study dedicated to the dynamics of muscle hyperplasia in several fish species concluded that fiber recruitment tends to cease when the body length reaches about half its maximum (7). However, due to the indeterminate growth of most fish species, this indication cannot be used to accurately estimate the length or weight at which hyperplasia ceases. Our thorough analysis of muscle fiber size distribution throughout the growth period of juvenile trout indicates that hyperplasia in white muscle decreases sharply up to 500 g, reaching a minimum plateau thereafter. This original result indicates that muscle hyperplasia declines much earlier than previously thought, but remains consistent with earlier work showing that total number of muscle fibers is reached around 1 to 2 kg (18) and that newly formed fibers are still present in white muscle of 1.2 kg trout (17). Indeed, the fiber size distribution shows the presence of small fibers up to 2 kg, but at very low frequency. In addition, the negligeable area (<0.5%) occupied by small fibers in fish weighing over 1 kg, shows that muscle growth is ensured almost exclusively by muscle fiber hypertrophy, with hyperplasia playing a marginal role.

The formation of new muscle fibers after hatching requires the activation and proliferation of SCs followed by differentiation and fusion of newly generated myocytes. Our thorough quantification of cells expressing the canonical SC marker Pax7 revealed a dramatic decrease in their numbers up to 500 g, followed by a minimal plateau up to 2 kg. This result is in agreement with our previous data showing that proliferative (BrdU^+^) cells in the SC niche (beneath basal lamina) were 3-fold less numerous in white muscle of 500 g than in 2 g trout (22). Thus, the decrease in SC number may result from their lower self-renewal rate. In Zebrafish, number of *pax7*^+^ cells decreases before hatching in white muscle and very rare positive cells are detected on adult (23, 24). In mice, different studies have reported an age-related decrease in the number of SCs (25–27) but differences have recently been observed between muscle types (28). Indeed, this number decreases with age in locomotor muscles (*Tibialis anterior* and *Extensor digitorum longus*) while it increases in the masseter and remains stable in other muscles. As the trout white muscle is a locomotor muscle, our results corroborate the idea that the SC number decline with aging like in equivalent vertebrate muscles.

In trout, our work reveals that the decrease in SC number parallels the decline in muscle hyperplasia, since both reach a minimal plateau in trout weighing 500 g, suggesting that the number of SCs is a key factor in the muscle hyperplasia decline. Nevertheless, in mouse *Extensor digitorum longus* muscle, SC number reaches a minimum plateau 14 days after the cessation of muscle hyperplasia around birth (2), indicating that other parameters are determining as previously suggested (29).

With aging and in the context of muscular dystrophies, it has been shown in mammals that myogenic capacities of SCs and their niche functionality are impaired (13, 14, 29).

To distinguish the specific role of SC intrinsic and extrinsic factors in the decline of muscle hyperplasia, transplantation of labeled donor cells into recipient tissue appeared to be the most relevant experimental approach. Our transplantation results indicate that only *mlc2*-GFP myogenic progenitors from early juvenile trout were able to differentiate and produce GFP^+^ myofibers, preserving their intrinsic myogenic capacity. Above a weight of 1 kg, our previous work has shown that SCs have low capacity for *in vitro* differentiation (22), even though they can still produce new muscle fibers at low rate in muscle of 1.5 kg trout (19). This decline in the intrinsic myogenic capacities of SCs observed in trout is reminiscent of that observed in mouse during aging (15, 29, 30). Our transplantation results also indicated that MDCs from 10 g *mlc2*-GFP trout produced GFP^+^ muscle fibers only in muscle of early juvenile trout recipients, which shows an alteration of the muscle niche functionality as early as late juvenile stage. Moreover, our results highlight that niche alteration that no longer allow the formation of new fibers occurs before the decrease in the intrinsic capacity of myogenic progenitors. These findings are in line with those obtained in rodent, revealing that aged tissue environment impairs myogenic capacities of young SCs to form new fibers and to regenerate muscle tissue (16, 31, 32). In the aged niche of mammals, several authors reported an increase in TGFβ signaling from fibers (31) associated with a decrease in Notch signaling (32) favoring activation and differentiation of SCs at the expense of their expansion. In addition, reduction of some component of the extracellular matrix such as fibronectin acts through p38αβ MAPK pathway to make SCs non-responsive to FGF2 signaling (33). It would be interesting in future studies to decipher these pathways in indeterminate growth models such as trout in order to understand whether the mechanisms involved could be comparable.

Taken together, the present study shows that myogenic capacities of SCs and the niche functionality are impaired in trout weighing over 500 g, in line with the low muscle regeneration capacities of 1 kg trout compared with 10 g. According to these results, we have already observed few formation of fibers during muscle regeneration of 1.5 kg trout in addition to a strong development of the connective tissue at the injury site (19), a typical landmark of aged muscle (34). In contrast to trout, adult zebrafish, which exhibit determinate growth, maintain good muscle regeneration capacity without significant connective tissue development (23). In line with these results, MDCs transplanted in adult zebrafish are still able to produced news fibers evoking that the niche has preserved its functionality (35). In fish with indeterminate growth, such as sea bream (*Sparus aurata*), almost complete regeneration of muscle has been reported without development of connective tissue (36–38). However, the fish used were at early juvenile stage (15-25 g) comparable to that of 10 g trout. These results suggest that fish with determinate growth retain a good capacity for muscle regeneration in adulthood, whereas those with indeterminate growth, such as trout or sea bream, see their muscle regeneration capacity declines after the hyperplastic growth period.

Of note, if the decline in myogenic capacity of SCs and niche functionality are hallmarks of aging in mammals, in return the juvenile trout of 500 g are young (1 year) and far from sexual maturation (2 years, 2-3 kg). Importantly, the comparable intrinsic capacity of myogenic progenitors observed in 12-month and 6-month-old trout of 20 g demonstrated that this decline is not due to aging but rather to increased weight. In other words, the period of exponential muscle growth up to 500 g induces a strong alteration of the intrinsic capacity of myogenic progenitors and also of the niche functionality, which is associated with a dramatic fall of SC number. During this period, the high rate of both muscle hyperplasia and hypertrophy requires extensive SC contribution, which may exhaust the pool of reserve cells and abolishes their self-renewal capacity. These results are reminiscent of those observed in the case of myopathies, where the chronic degeneration processes lead to the SC depletion (14). In contrast, it is interesting to note that fish with determinate growth such as zebrafish do not undergo muscle hyperplasia after hatching, thus preserving the SC pool and possibly explaining the efficient muscle regeneration observed in adult. In line with these observations, it has been shown that the number of MDCs extracted from adult fish is inversely related to the muscle mass, with trout having the lowest quantity (39). For example, *Danio dangila*, a specie closely related to zebrafish (*Danio rerio*) but with indeterminate growth, has 6 times fewer MDCs than zebrafish.

In conclusion, our study shows that the arrest of muscle hyperplasia in juvenile trout results from an exhaustion of SCs as evidenced by a rapid fall of their pool and their intrinsic capacity, but also from a loss of functionality of their niche. These features, present in some myopathies and muscle aging, may also partly explain the arrest of hyperplasia around birth in mammals and birds. This work underscores the relevance of trout, whose evolution of muscle hyperplasia is highly dynamic, as a model for studying muscle biology and highlights the importance of intrinsic SC properties and niche environment in the formation of new muscle fibers in adulthood.

## Materials and Methods

### Animals

Immature rainbow trout (*Oncorhynchus mykiss*) were reared in a recirculating rearing system under natural simulated photoperiod and at 12 ± 1 °C. Fish were fed daily *ad libitum* on a commercial diet. The *mlc2*-GFP transgenic trout line was produced to express GFP specifically in the differentiated muscle cells (21). All the experiments were developed in accordance with the current legislation governing the ethical treatment and care of experimental animals (order no. 2001-464; European directive 2010/63/EU). They were conducted at the INRAE Fish Physiology and Genomic Laboratory (LPGP) experimental facilities (DOI:10.15454/45d2-bn67, permit number D35-238-6, Rennes, France) and approved by the ethics committee for the animal experimentation of Rennes and the French minister of national education, research, and innovation, under the authorization number APAFIS# 10496-2017051815207923, #35543-202202181607249 and # 44261-2023071915256688.

### Production of 20 g and 1 kg trout of the same age

As a poikilotherm with continuous growth, trout exhibit extraordinary muscle growth plasticity (40). To obtain 20 g and 1 kg fish at the same age, trout from the same fertilization were fed from the first-feeding stage either 1-2 days/weeks or fed manually *ad libitum* during 11 months. Three weeks prior to transplantation the restricted trout were fed daily to ensure full growth.

### Muscle-derived cell extraction

For all studies, MDCs were isolated from the dorsal part of the white muscle of transgenic *mlc2*-GFP trout as previously described (41). Briefly, 10 to 80 g of muscle were collected in cold Dulbecco’s modified Eagle’s medium (DMEM; #D7777, Sigma) supplemented with 9 mM NaHCO3, 20 mM HEPES, 15% horse serum, and antibiotics (100 U/ml penicillin, 100 μg/ml streptomycin). Muscle was mechanically dissociated and enzymatically digested by collagenase (#C9891, Sigma) and trypsin (#4799, Sigma) following by successive 100 μm- and 40 μm-filtrations. Cells were resuspended in DMEM containing 10% fetal bovine serum (FCS) and 1% antibiotic-antimycotic solution and counted.

### Muscle-derived cell transplantation

Freshly extracted MDCs were maintained on ice until transplantation (<1 h). To easily locate the transplantation area, 1% of fluorescent red tracer (#F8826, Molecular Probes) was added to the cell suspension. The concentration of the cell suspension was adjusted to 4×10^6^ cells/ml to inject 1×10^5^ cells to each fish (25 µl). Prior to transplantation, recipient trout were anesthetized with MS-222 at 50 mg/l and placed on a damp cloth. Using a sterile 0.45 mm needle, the cells were transplanted into dorsal white muscle (4 mm depth) behind the dorsal fin and above the lateral line. Three weeks later, fish were euthanized by an overdose of MS-222 (200 mg/l) and muscle samples from transplanted area were taken and split into two parts. Then, both parts were imaged in red (tracer) and green (GFP) fluorescence with a Nikon Multizoom microscope (AZ100). For each fish, control muscle sample from the contralateral part was collected. Finally, muscle samples were fixed for 2 hours in 4% paraformaldehyde (PFA) on ice, dehydrated and embedded in low-melting polyester wax (#19312, Electron microscopy Science).

### Muscle histology

Muscle samples were collected from 10 g (11 g; 11 cm), 100 g (141 g; 20 cm), 500 g (542 g; 31 cm), 1 kg (1088 g; 37 cm), 1.5 kg (1534 g; 41 cm), and 2 kg (2083 g; 44 cm) trout and fixed in 4% PFA overnight at 4°C and embedded in paraffin. Cross-sections were then cut at 7 µm using a microtome (HM355; Microm Microtech, Francheville, France), deparaffinized, rehydrated and stained with Alexa 488-conjugated wheat germ agglutinin (5 µg/ml in PBS; WGA; Molecular Probes # W11261) for 3 hours at room temperature, to visualize connective tissue and basal lamina (42). Samples of transplanted muscle were deparaffinized and rehydrated using serial dilution of ethanol and then permeabilized 3 min in 0.1% Triton X-100/PBS. After three washes, sections were saturated for 1 h with 3% bovine serum albumin, 0.1% Tween 20 in PBS (PBST). Sections were incubated for 3 h with myosin antibody (#MF20, Hybridoma Bank) diluted in blocking buffer. The anti-mouse Alexa 350-conjugated antibody (#A11045, ThermoFisher) was diluted in PBST and applied for 1 h. Sections were mounted with Mowiol containing 4,6-diamidino-2- phenylindole (DAPI; 0.5 μg/ml). All the images were taken with a Nikon digital camera coupled to a Nikon 90i widefield fluorescence microscope.

### In situ hybridization

*In situ* hybridization was performed on 7 µm-muscle cross-sections using the chromogenic RNAscope® 2.5HD detection reagent RED kit (#322360; Bio-Techne) according to the manufacturer’s protocol. Briefly, sections were briefly baked at 60°C for 1 h, deparaffinized and air dried. After 10 min in hydrogen peroxide solution (#322335, Bio-Techne), sections were treated with 1X Target Retrieval (#322000; Bio-Techne ) for 15 min at 100°C, followed by 25 min with Protease Plus solution (#322331; Bio-Techne) at 40°C. Due to the presence of three pax7 genes in the trout genome (43), a set of probes able to recognized *pax7a1*, *pax7a2* and *pax7b* mRNA was designed (20). This set was hybridized for 2 h at 40°C. All steps at 40°C were performed in a Bio-Techne HybEZ II hybridization system (#321720). Nuclei were counterstained with 0.5 µg/ml DAPI and sections were mounted with EcoMount (#EM897L, Biocare medical). In the chromogenic RNAscope assay, the red signal (Fast Red) can be observed with either a white light or fluorescence microscope.

### Automated analysis of images

Fiber size distribution as a function of fish weight was determined on 5 images of the white muscle transversal section for each fish (10–12). Cellpose 2 (44) was first used to detect muscle fibers, followed by Fiji (45) to measure the surface area of each fiber. Muscle fiber diameters (D) were then calculated using the formula D = 2√(surface/π), assuming that the cross-sections of individual fibers were circular.

Quantification of *pax7*^+^ cells was also performed on these images. All DAPI-labelled nuclei were detected with the Fiji stardist plugin (46), then nuclei with an intensity in the red channel greater than 100 were considered *pax7* positive.

The distribution of GFP^+^ and GFP^−^ fibers was obtained using Visilog 6.8 software. Muscle fibers were first detected on the basis of blue channel intensity, then manually corrected in case of erroneous detection. GFP^+^ fiber was determined as a fiber containing more than 50% green pixels after a thresholding above 30 gray values on the green channel.

The GFP signal on transplanted area was quantified using a macro-command on Fiji software. Basically, the green (GFP) and red (tracer) area above a fixed threshold were quantify and the ratio GFP/Red area was calculated.

### Statistical analyses

Data were analyzed using the non-parametric Kruskal-Wallis rank test followed by the *post-hoc* Dunn test. All analyses were performed with the R statistical package (version 4.3.2).

## Acknowledgments

This work was supported by the ANR FishMuSC (ANR-20-CE20-0013-01). The authors thank Lucas Génevé and Alex Choplin for their help in measuring *pax7*^+^ cell density.

---

**Supplemental Figure S1:**
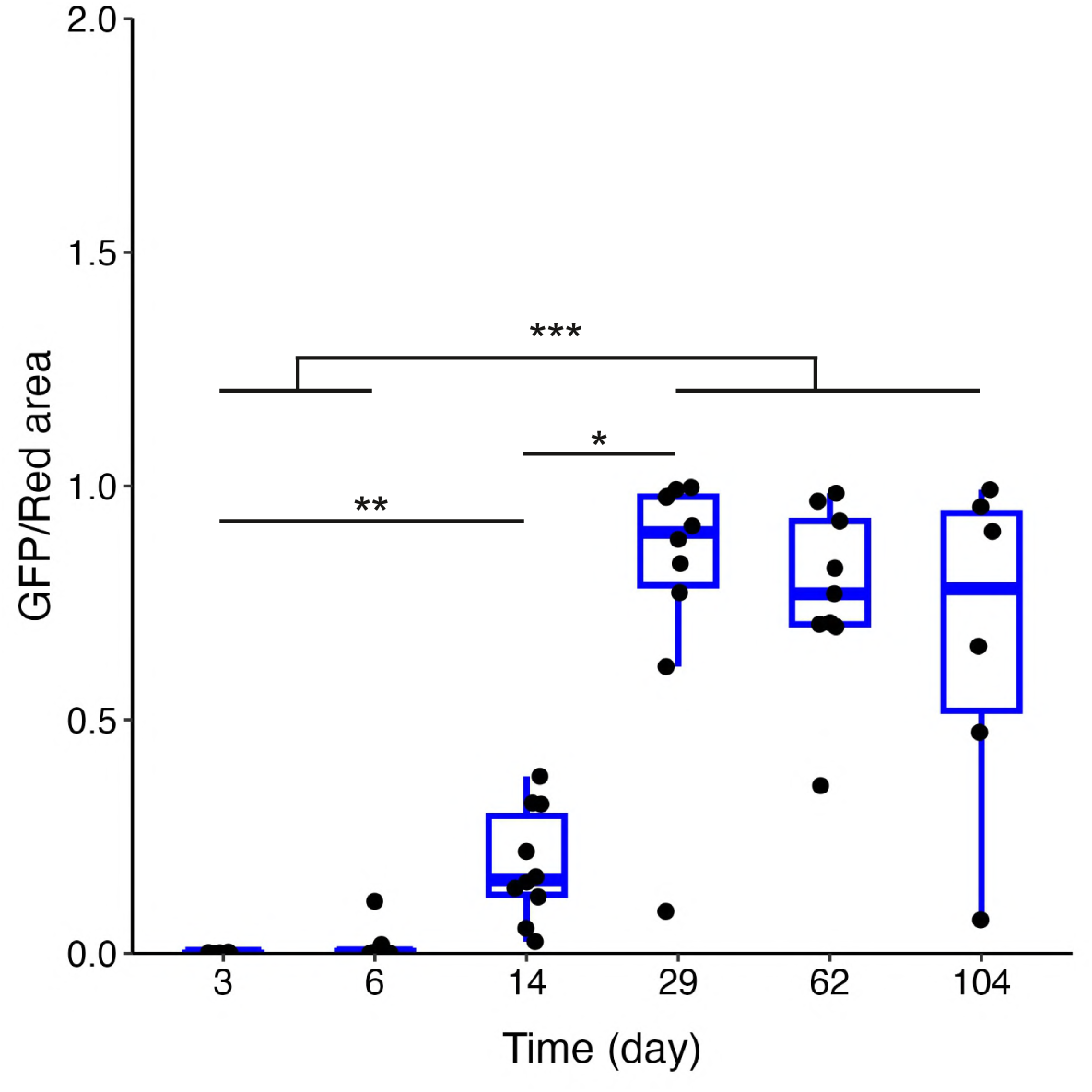
*mlc2*-GFP muscle-derived cells transplanted into wild-type trout were able to differentiate. *Mlc2*-GFP muscle stem cells from 10 g trout were transplanted into muscle of 10 g wild-type trout. For each time point, muscle samples from the transplantation site were collected and pictures were taken to measure red tracer and GFP signal surface using a Nikon multizoom macroscope. Data present the ratio between the GFP and the red surface. (N = 6-10). All statistical significances were calculated by the non-parametric Kruskal–Wallis rank test followed by the post-hoc Dunn test. Significance levels: *p<0.05, **p<0.01, ***p<0.001.

**Supplemental Figure S2:**
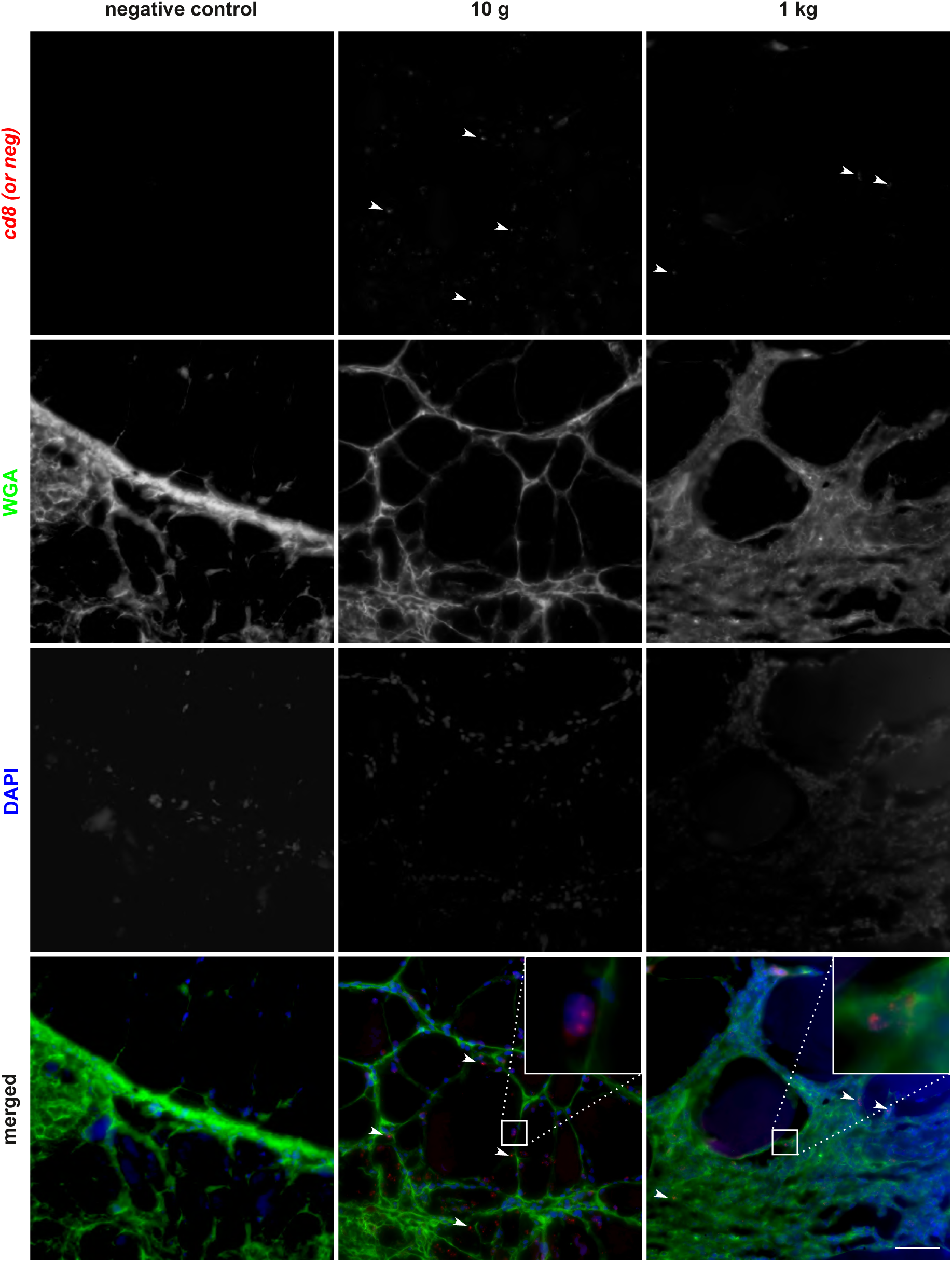
Transplantation of muscle-derived cells did not induce an immune rejection reaction at the injection site. Cross-sections of trout white muscle were performed 21 days after transplantation and analyzed by *in situ* hybridization for *cd8* (red, marker of T-cells) and the extracellular matrix was stained with Alexa 488-conjugated wheat germ agglutinin (WGA; green). No signal was observed with the negative probe against a bacterial gene *DapB.* The nuclei are counterstained with DAPI and the scale bar corresponds to 100 µm.

**Supplemental Figure S3:**
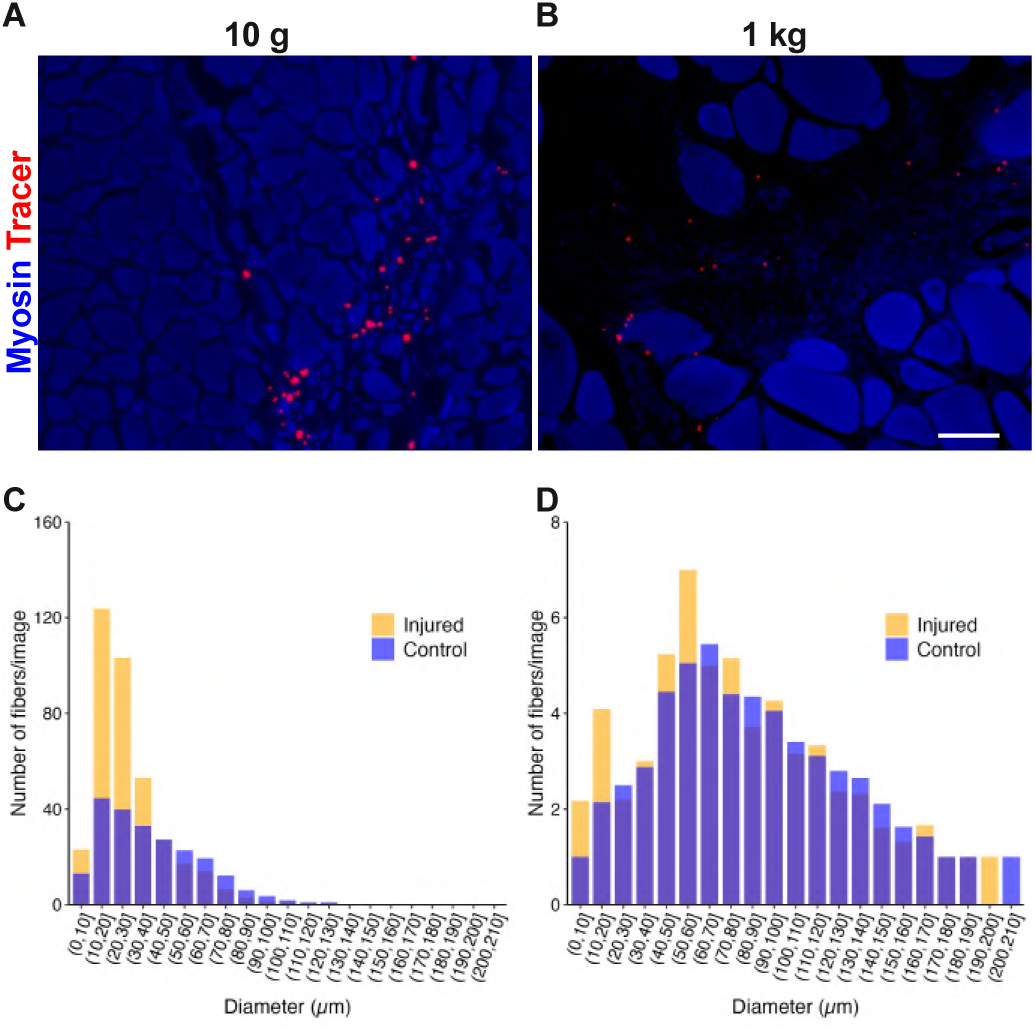
Muscle regeneration potential declines early during the juvenile period in trout. Pictures of muscle cross-sections showing immunolabeling for myosin (blue) and tracer (red) 21 days after injury (A, B). The scale bar corresponds to 100 µm. Panels A and C, correspond to injury of 10 g trout muscle. Panels B and D, correspond to injury of 1 kg trout muscle. Panels C and D correspond to fiber size distribution in control and injured muscle. (N = 9-10, except for 10 g injured fish where only 4 fish could be analyzed). To assess the combined effects of trout weight and muscle condition (healthy vs. injured) on fiber diameter distribution, a generalized linear mixed model (GLMM) with a log-normal distribution was applied. The dependent variable was the log-transformed adjusted fiber diameters (log(diameter + 1)). The fixed effects included muscle condition, trout weight (10 g vs. 1 kg), and their interaction (p<0.001 for each). A random intercept for each image was also included to account for variability between images. Model fitting and statistical significance testing were performed using the glmmTMB package (version 1.1.9) in R (version 4.3.2). Residual diagnostics were conducted to ensure model assumptions were met.

